# Detection of antimicrobial resistance, pathogenicity, and virulence potentials of non-typhoidal *Salmonella* isolates at the Yaounde abattoir using whole genome sequencing technique

**DOI:** 10.1101/2021.12.15.472740

**Authors:** Chelea Matchawe, Eunice M. Machuka, Martina Kyalo, Patrice Bonny, Nkeunen Gerard, Isaac Njaci, Seraphine Nkie Esemu, Dedan Githae, John Juma, Mohamadou Bawe, Bonglaisin J. Nsawir, Edi Piasentier, Lucy M. Ndip, Roger Pelle

## Abstract

One of the crucial public health problems today is emerging and re-emerging of multidrug-resistant bacterial pathogens coupled with a decline in the development of new antimicrobials. Non-typhoidal *Salmonella* is classified among the multidrug-resistant bacterial pathogens of international concern. To predict their multidrug resistance potentials, 19 assembled genomes (partial genomes) of 23 non-typhoidal *Salmonella* isolated at the Yaounde abattoir between December 2014 and November 2015 from live cattle (n=1), beef carcass (n=19), butchers’ hands (n=1) and the beef processing environments (n=2) were explored using whole-genome sequencing. Phenotypically, while approximately 22% (n=5) of *Salmonella* isolates showed moderate resistance to streptomycin, 13.04 % (n=3) were multidrug-resistant. Genotypically, all the *Salmonella* isolates possessed high multidrug resistance potentials against several classes of antibiotics (third-generation cephalosporin and fluoroquinolone), which are assigned highest priority drugs by the World Health Organization. Moreover, more than 31% of the isolates exhibited resistance potentials to polymyxin, considered as the last resort drug with both clinical and veterinary relevance. Additionally, close to 80% of non-typhoidal *Salmonella* isolates in this study harboured ‘‘silent resistant genes’’ and thus constituted potential reservoirs of antibiotic resistance to other foodborne bacteria. Plasmids also appear to play a critical role in the horizontal transfer of antibiotic resistance genes of some isolates. The isolates showed a high degree of pathogenicity and possessed key effector proteins to establish infection in their hosts, including humans. The overall results demand prudent use of antibiotics and constant monitoring of antimicrobial resistance of non-typhoidal *Salmonella* in the Cameroonian abattoirs.

**Author summary:** Non-typhoidal *Salmonella* has been classified among the multidrug resistant bacterial pathogens of international concern. A growing resistance to a broad range of antibacterial compounds in animals and clinical settings has been reported in Non-Typhoidal *Salmonella*. Current knowledge of their antibiotic resistance profile is essential to inform policy decisions for the choice of appropriate management of invasive salmonellosis. The significance of our research consists in predicting the multidrug resistance, pathogenicity and virulence potentials of *Salmonella* organisms using whole genome sequencing. This unveils the need for the development of a diagnostic model that takes into account the genotype–phenotype antibacterial resistance profile of *Salmonella*, which is of both clinical and veterinary relevance.

## Introduction

Today’s world is experiencing an antibiotic resistance pandemic due to a growing bacterial resistance to a broad range of drugs in animals and clinical settings [1,2,3]. Microbial multidrug resistance (MDR) frustrates efforts for infection control resulting into a considerable increase in morbidity and mortality worldwide [1, 2].

MDR has also been reported in non-typhoidal *Salmonella* (NTS). For instance, while 16% of NTS isolates exhibited resistance to at least one essential antibiotic, as high as 2% of them were resistant to at least three different classes of antibiotics in the US [2]. In Europe, 23% of NTS isolated from calf carcasses were MDR [3]. As a result, WHO considered NTS as a priority pathogen, thereby ranking them among the MDR bacterial pathogens of international concern [1]. Animals, including cattle, are the main sources of antibiotic-resistant bacteria [3] owing to the use of antibiotics for disease prevention or as animal growth promoter.

Current knowledge of the type of *Salmonella* serovars and their antibiotic resistance profile is essential to inform policy and guide treatment strategies for appropriate therapy and development of new antimicrobials [1, 2]; thus WHO recommendation for national surveillance of antimicrobial resistance (AMR) in *Salmonella* [1]. Given their zoonotic nature, there is a need for an integrative ‘One Health’ approach for the surveillance of AMR among humans and animal *Salmonella* isolates [4]. Unlike traditional antimicrobial susceptibility testing, whole-genome sequencing (WGS) can give information on the presence of MDR genes [5] and pathogenicity and virulence factors in *Salmonella* organisms. This study was aimed at predicting MDR, pathogenicity, and virulence potentials of *Salmonella* isolated at the Yaounde abattoir using WGS.

## Results

### Phenotypic antibiotic resistance profile of *Salmonella* isolates

Twenty-three genomes were successfully sequenced out of 38 *Salmonella* isolates identified. However, only 19 sequenced genomes were thoroughly exploitable for all the required bioinformatics analyses.

Isolates **8ev**, **20de**, **22sa**, **34de**, **60sa**, and **evjul** were resistant to streptomycin, whereas between 18 and 20 isolates were highly susceptible to ampicillin, chloramphenicol, and tetracycline (Table 1). However, isolates **8ev**, **22sa** and **34de** were multidrug resistant (MDR).

**Table 1:** Antimicrobial sensitivity of non-typhoidal *Salmonella* isolated at the Yaounde abattoir.

| Sample code | *Salmonella* isolate | Tetracycline (Ø= mm) | Chloramphenicol (Ø= mm) | Streptomycin (Ø= mm) | Ampicillin (Ø= mm) |
| --- | --- | --- | --- | --- | --- |
| 34ev | Wernigerode | 30 | 34 | 24 | 22 |
| 8ev | Poona | 9 (≤ 11) | 7 (≤ 12) | 6 (≤ 11) | 6 (≤ 13) |
| 35dea | Wilhelmsburg | 28 | 35 | 25 | 22 |
| 35deb | Wilhelmsburg | 30 | 31 | 24 | 20 |
| 22sa | Poona | 10 (≤ 11) | 6 (≤ 12) | 6 (≤ 11) | 6 (≤ 13) |
| 31eva | Wilhelmsburg | 33 | 40 | 25 | 22 |
| 31evb | Wilhelmsburg | 38 | 46 | 25 | 20 |
| 32eva | Wilhelmsburg | 35 | 40 | 25 | 25 |
| 32evb | Wernigerode | 30 | 31 | 25 | 23 |
| 86ev | Infantis | 27 | 35 | 8 | 14 (14-16) |
| 100ev | Wernigerode | 32 | 40 | 24 | 22 |
| 36ev | Wilhelmsburg | 29 | 38 | 25 | 22 |
| 98se | Wernigerode | 32 | 40 | 26 | 18 |
| 108ev | Kibusi | 30 | 38 | 25 | 17 |
| 20de | Enteritidis | 32 | 35 | 8 (≤ 11) | 22 |
| 34de | Poona | 10 (≤ 11) | 10 (≤ 12) | 6 (≤ 11) | 7 (≤ 13) |
| 60sa | Enteritidis | 34 | 35 | 13 (12-14) | 29 |
| 88sa | Infantis | 32 | 40 | 25 | 24 |
| 88sab | Infantis | 33 | 38 | 24 | 22 |
| 133sa | Enteritidis | 32 | 36 | 25 | 14 (14-16) |
| 103bo | Wilhelmsburg | 34 | 40 | 24 | 20 |
| EVJUL | Mbandaka | 28 | 33 | 9 (≤ 11) | 18 |
| DEF1 | Not sequenced | 30 | 36 | 22 | 20 |

NB: Zone of inhibition in red indicates resistance to the test antibiotic; blue = intermediate; in black are the numbers representing the diameter of zone of inhibition mean susceptible to the test antibiotics; code of *Salmonella* strain written in red, represent the multidrug-resistant (MDR) strains.

### Distribution of resistance genes against specific antibiotic across *Salmonella* isolates

The distribution of specific antibiotic resistance genes across the *Salmonella* isolates studied is summarized in Table 2. Genes *aadA, aadA1*, and *aadA2* conferring specific resistance to streptomycin were respectively present in 5, 21, and 10.5% of the *Salmonella*. Moreover, chloramphenicol-resistance was found in more than 15% of the isolates possessing *cat1, cat2*, and *cat3* genes while 10.5% of the isolates haboured *cmlA1, cml5*, and *cml6* genes. The genes *tet(A), tet(B), tet(C)* and *tetR* genes conferring resistance to tetracycline were present in 5.26% 10.52%, 10.52%, 15.78% and 10.52% of the isolates, respectively. TEM-1 and TEM-163 genes conferring resistance against ampicillin were identified in 5.26% and 10.52% of *Salmonella* isolates, respectively.

**Table 2:**
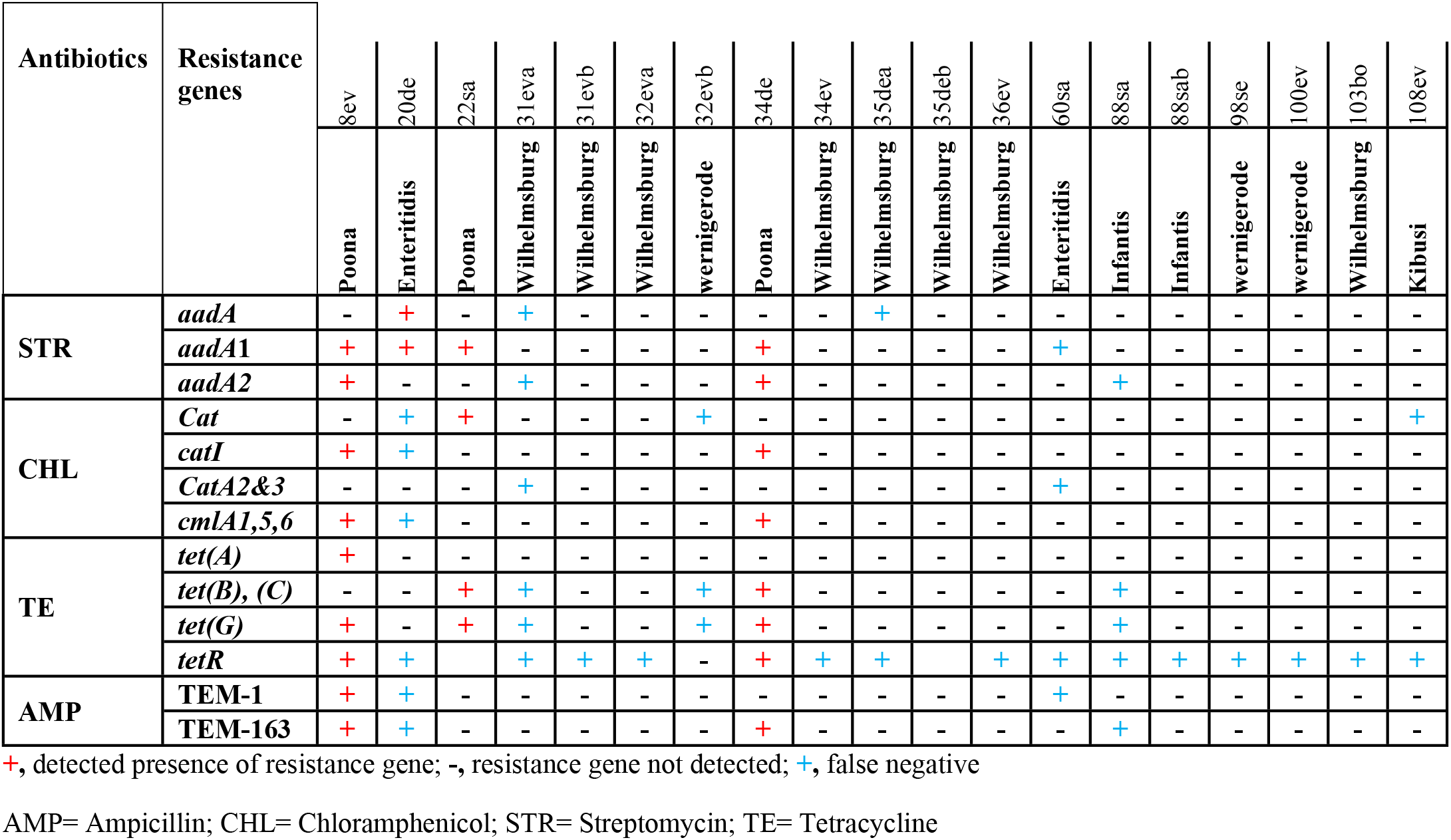
Distribution of resistance genes against specific antibiotic across *Salmonella* isolates in the study.

However, false-negative results or “silent resistance genes” were reported (Table 2). Close to 80% of *Salmonella* isolates harbored at least one “silent gene”. These genes were detected in the susceptible *Salmonella* isolates but failed to be expressed phenotypically.

### Multidrug resistance potential of *Salmonella* serovars

Eighteen genes (in purple color) involved in the efflux, transport, and reduced permeability of antimicrobials conferring thereby multidrug resistance to *Salmonella* were identified (Table 3). The presence of *TolC* was reported in all the *Salmonella* isolates but was only found in 5.26% isolates for the macrolides. The *golS* gene, a promoter for MdsABC complex, a multidrug efflux pump, was detected in all the isolates (n = 19). Interestingly, the MdsABC (multidrug transporter of *Salmonella*) complex comprising *mdsA*, *mdsB*, and *mdsC* units was also present alongside with gene *golS* in 100% isolates that exhibited potential resistance to phenicols. The *E.coli soxS* and *soxR* genes were detected in 78.94 to 100% *Salmonella* isolates.

**Table 3:**
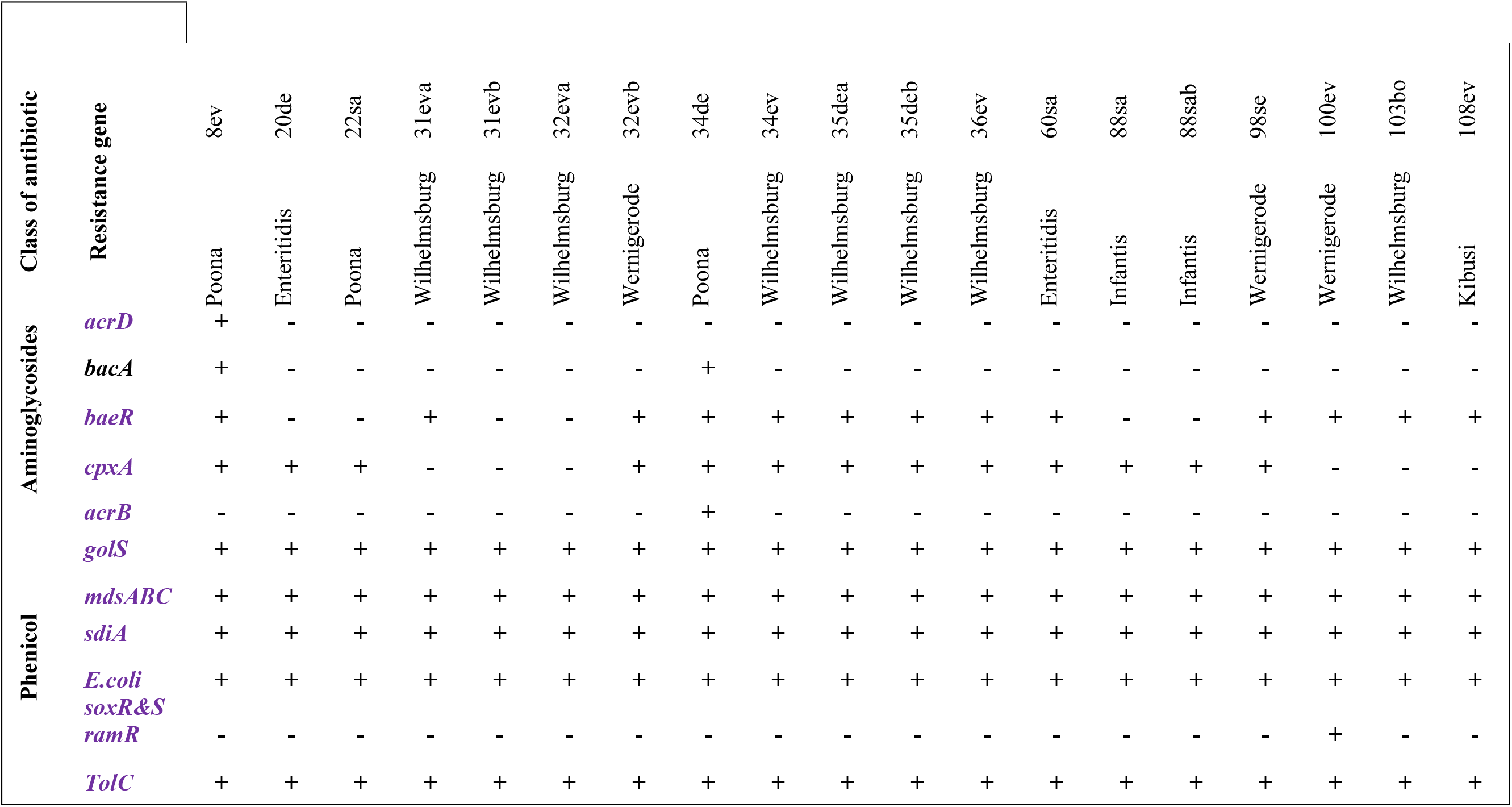
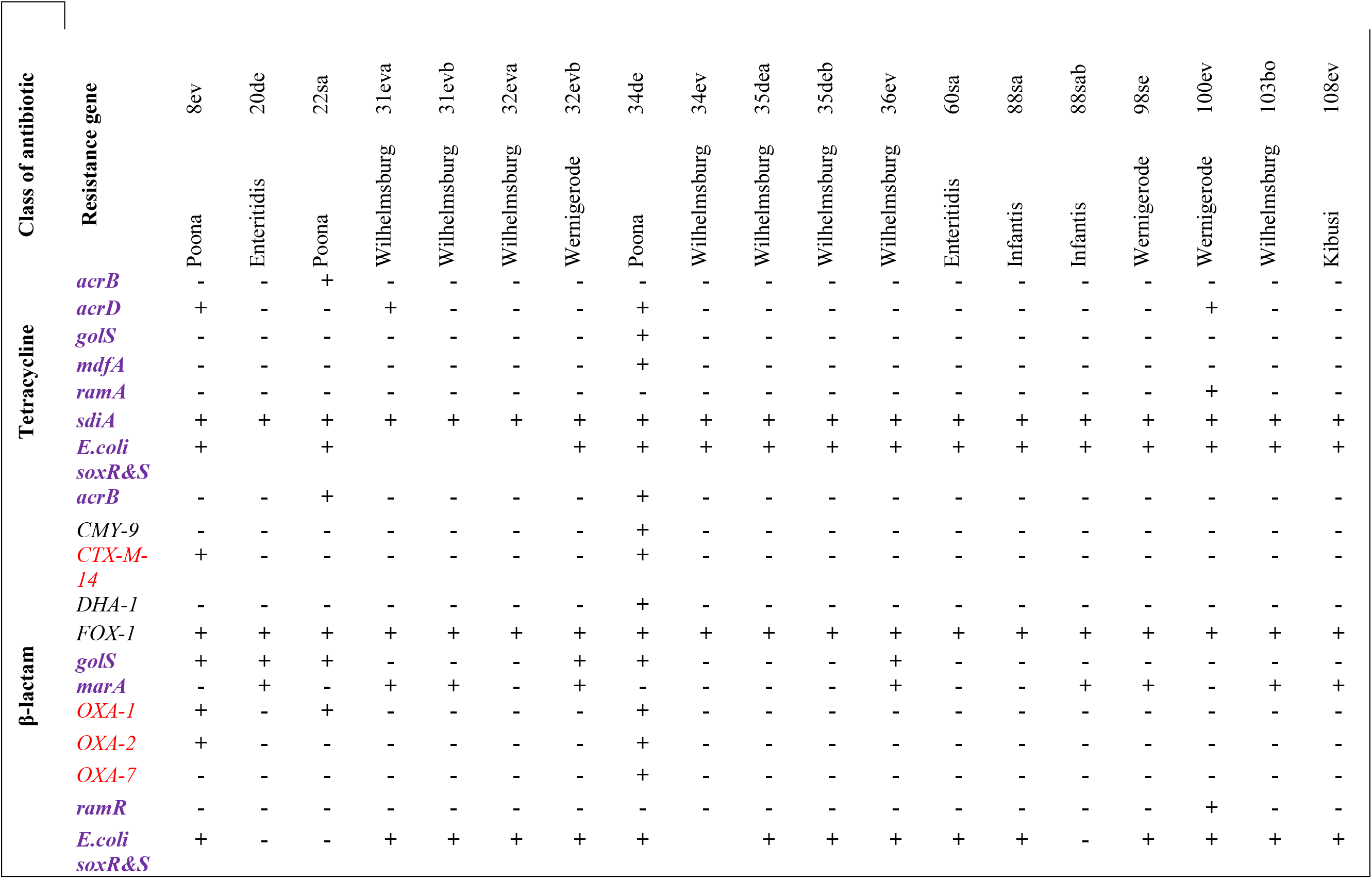
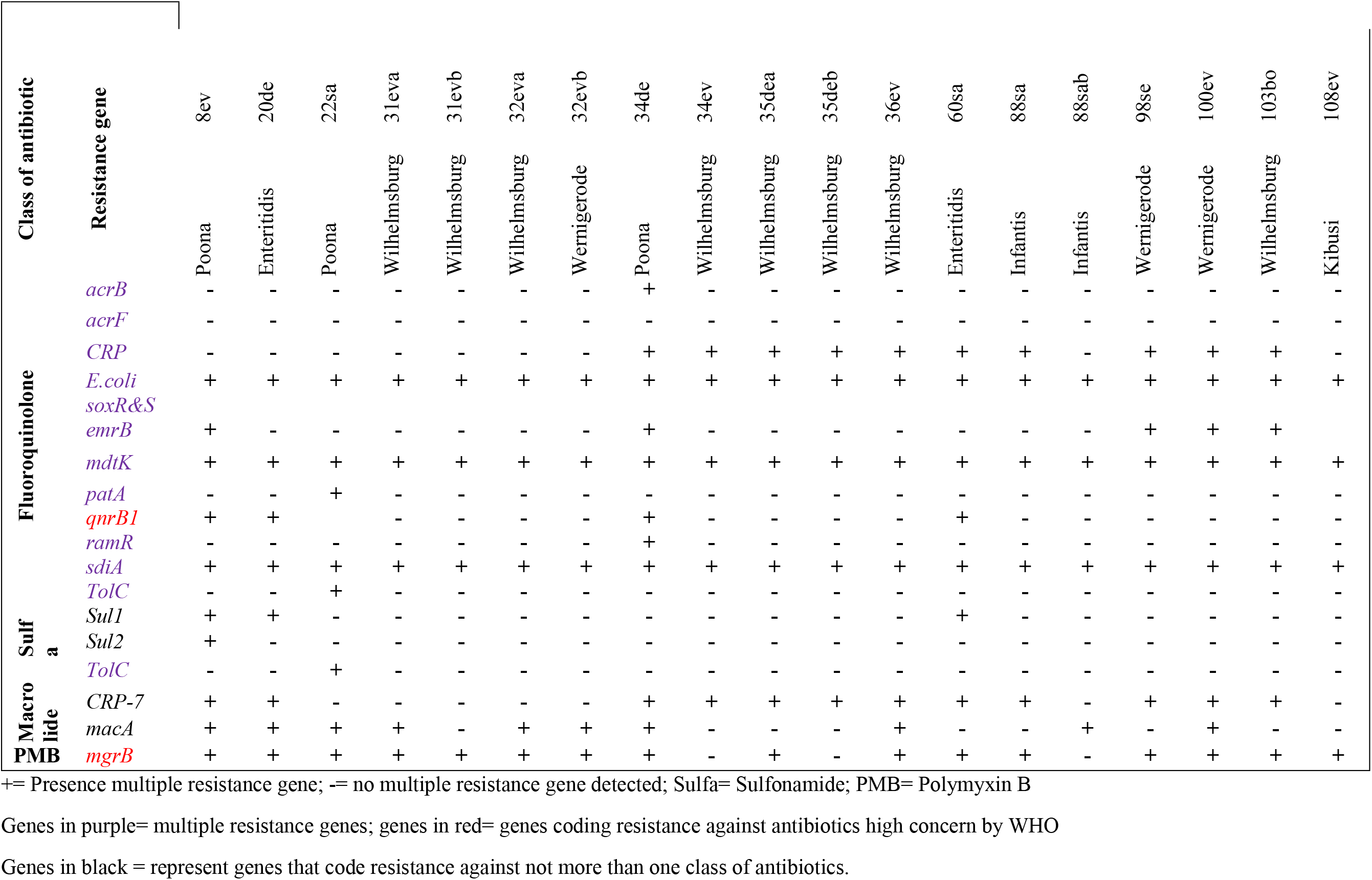
Distribution of multidrug resistance genes among *Salmonella* serotypes.

On the other hand, the gene *mdtk*, a multi-efflux pump conferring resistance mainly against fluoroquinolone, was present in 100% *Salmonella* isolates. The gene *sdiA*, a regulator for multi-drug resistance pump *AcraB*, was detected in 100% of the isolates.

Besides, other genes such as *sul1* and *sul2* that confer specific resistance against sulfonamides and *CTX-M-14* that encodes for resistance against 3^rd^ generation cephalosporin were also noted in 15.8 and 5.26% *Salmonella* isolates, respectively. Three *Salmonella* isolates (**8ev, 22sa** and **34de**) harbored genes *OXA (OXA-1, OXA-2* and *OXA-7*) conferring resistance against cephalosporins. In addition, *qnrB*1 gene that confers resistance against fluoroquinolone was reported in isolates **8ev**, **20de**, **34de** and **60sa**.

The gene *macA* that mediates efflux of macrolide antibiotics and secretion of enterotoxin ST11 via AcrAB-TolC efflux pump was detected in 52.63% NTS isolates. Additionally, the gene *marA*, which mediates efflux of several antibiotics and disinfectants via AcrAB-TolC efflux pump, was expressed by 47.36% *Salmonella* isolates.

### Polymorphism-induced potential resistant effect against Polymyxins of *Salmonella* serovars

Mutations on the PmrA/PmrB genes (Table 4) conferring resistance potential against polymyxins were detected in more than 31% isolates.

**Table 4:** Polymorphism in *Salmonella* serovars with potential resistance to polymyxins.

| Isolate code | Position of mutation on pmrA & pmrB | Nucleotide change | Amino acid change |
| --- | --- | --- | --- |
| **8ev** | **pmrB** |  |  |
|  | pmrB p.M15T | ATG → ACT | M → T |
|  | pmrB p.G73S | GGC → AGC | G → S |
|  | pmrB p.V74I | GTA → ATA | V → I |
|  | **pmrA** |  |  |
|  | pmrA p.T89S | ACC → AGC | T → S |
| **22sa** | **pmrB** |  |  |
|  | pmrB p.M15T | ATG → ACT | M → T |
|  | pmrB p.G73S | GC → AGC | G → S |
|  | pmrB p.A111T | GCG → ACG | A → T |
|  | **pmrA** |  |  |
|  | pmrA p.T89S | ACC → AGC | T → S |
| **31eva** | **pmrB** |  |  |
|  | pmrB p.M15T | ATG→ACT | M → T |
|  | pmrB p.G73S | GGC → AGC | G → S |
|  | pmrB p.V74I | GTA → ATA | V → I |
|  | pmrB p.A111T | GCG → ACG | A → T |
|  | pmrA p.T89S | ACC → AGC | T → S |
| **32eva** | **pmrB** |  |  |
|  | pmrB p.I18L | ATT→CTT | I → L |
| **34de** | **pmrB** |  |  |
|  | pmrB p.M15T | ATG → ACT | M → T |
|  | pmrB p.G73S | GGC → AGC | G → S |
|  | pmrB p.V74I | GTA → ATA | V → I |
|  | pmrB p.A111T | GCG → ACG | A → T |
|  | pmrB p.L352M | CTG → ATG | L → M |
|  | **pmrA** |  |  |
|  | pmrA p.T89S | ACC → AGC | T → S |
| **35dea** | **pmrB** |  |  |
|  | pmrB p.V126G | GTC → GGC | V → G |
|  | pmrB p.S127A | TCG → GCG | S → A |
|  | pmrB p.I129L | ATC → CTC | I → L |
|  | pmrB p.V133D | GTT → GAT | V → D |
|  | pmrB p.L136R | TTG → AGG | L → R |
|  | pmrB p.T139P | ACG → CCG | T → P |

### Pathogenicity and virulence factors

All the *Salmonella* isolates had high probability (with a mean probability of 94%) to cause diseases in humans (Table 5). Moreover, the proteome of the *Salmonella* isolates has matched with a broad range of pathogenic bacterial families (466-787). Serovar Enteritidis had the highest prediction probability (p=0.95) as human pathogen while serovar Poona registered the lowest (p=0.93).

**Table 5:** Prediction of *Salmonella* isolates as human pathogens.

| Serovars | PHPathogen | Proteome coverage (%) | Matched PF | Non PF |
| --- | --- | --- | --- | --- |
| Poona | 0.93 | 10.1 | 466 | 5 |
| Enteritidis | 0.95 | 18.2 | 787 | 2 |
| Wilhelmsburg | 0.941 | 15.71 | 691 | 3 |
| Wernigerode | 0.944 | 15 | 661 | 3 |
| Infantis | 0.941 | 17.45 | 746 | 3 |
| Kibusi | 0.943 | 18 | 778 | 3 |

Key: PHPathogen = predicted as human pathogen; PF = pathogenic family; NPF= Non-pathogenic family

Eleven SPIs including C63PI have been identified as outlined in Table 6. Only C63PI was present in all the isolates. The function of each virulence factor is summarized in Table 7. Other virulence factors including effector proteins, adhesion factors, virulence plasmids, and toxins were also detected in our isolates (Tables 8a and 8b).

**Table 6:**
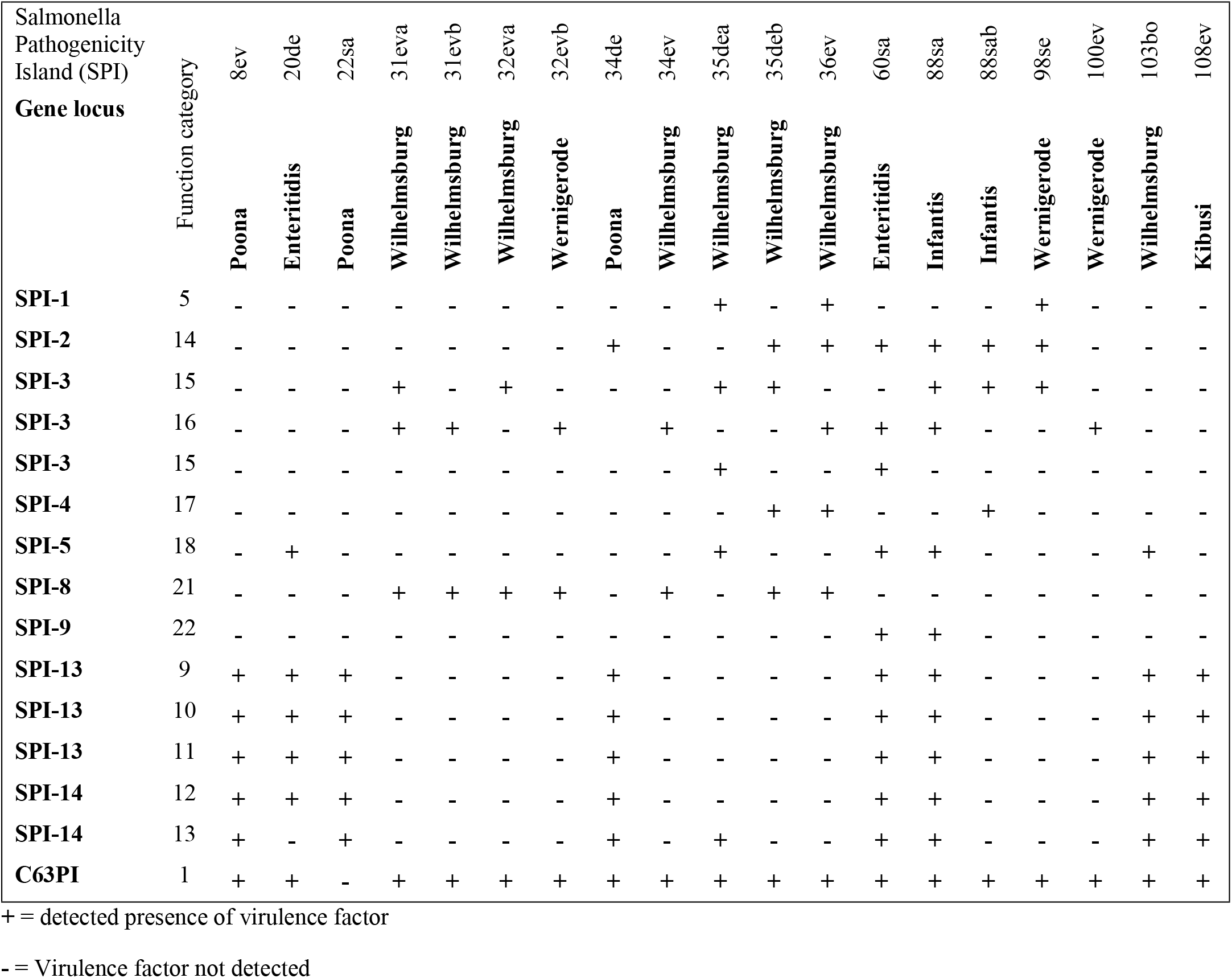
virulence factors among Salmonella serovars.

**Table 7:** Salmonella pathogenicity island-encoded effector proteins.

| Isolates | Protein effector | Function | Origin |
| --- | --- | --- | --- |
| 8ev, 20de, 31eva, 31evb, 32eva, 32evb 34de, 34ev, 35dea, 35deb, 36ev, 60sa, 88sa, 88sab, 98se, 100ev, 103bo, 108ev | Cell adherence/invasion proteins : SipA, EnvF, InvA, InvE, SipB, SipC, SipD, SopB, SopE, Sii E, IagB, SptP, protein oxygen-regulated invasion protein OrgB, PhoP, PhoH protein, protein MisL, host colonization factor (ShdA), protein rtn T 6SS protein IcmF | involved in adherence, cell aggregation, biofilm formation, cell invasion, resistance to antimicrobial peptides, internalization and survival within macrophages | Enteritidis P125109, Paratyphi A ATCC 9150, Newport SL254, Agona SL483, Typhimurium LT2, Schwarzengrund Choleraesuis SC-B67Heidelberg SL476 |
| 31eva, 31evb, 32eva, 34de, 34ev, 36ev, 88sab, 98se, 100ev, 103bo | OmpL | Outer membrane protein with high immunogenicity potential | Schwarzengrund, Newport SL254 |
| 20de, 22sa, 31evb, 32evb, 36ev, 60sa, 88sab | Heat shock protein, Stress proteins | Small polypeptides that play important role during *Salmonella* pathogenesis | Choleraesuis SC-B67, Klebsiella pneumoniae |
| 8ev, 20de, 31eva, 31evb, 32eva, 34de, 34ev, 35dea, 36ev, 60sa, 88sa, 88sab, 98se, 100ev, 103bo, 108ev | T3SS effector protein SopA, SopE, secreted effector protein SopE2, virulence proteins SopD and SopD2, Effector proteins pipB2, SpiC, SpvB, SseB, SseC, and SseD, T6SS lysozyme-related protein, Ais protein, effector proteins IpaD/SipD, protein bax | Promote *Salmonella* survival replication within macrophages, critical for virulence in different hosts, in invasion of epithelial cells and intestinal inflammation, translocation of SPI-1, SPI2 effector proteins<br>Involved in proapoptotic activity | Enteritidis P125109, Paratyphi A ATCC 9150, Newport SL254, Paratyphi C RKS4594 Schwarzengrund, Agona SL483 Heidelberg SL476 |
| 20de, 31eva, 31evb, 32eva, 34de, 34ev, 36ev, 60sa, 88sa, 88sab, 98se, 100ev, 108ev | Antigen presentation protein SpaK, SpaN, SpaS, SpaR, | Responsible for the surface presentation of determinants needed for the entry of *Salmonella* species into host cells | Agona SL483, Typhimurium LT2, Newport SL254r. SL483 |
| 8ev, 20de, 31eva, 31evb, 32eva, 32evb, 34de, 34ev, 36ev, 60sa, 88sa, 88sab, 98se, 103bo, 108ev | EsrB, secretion system apparatus SsaV, SsaH and SsaM, SPI-6 encoded Rhs-protein | Response regulators of secreted effectors and host inflammatory responses | Schwarzengrund CVM19633, Newport SL254 |

**Table 8a:**
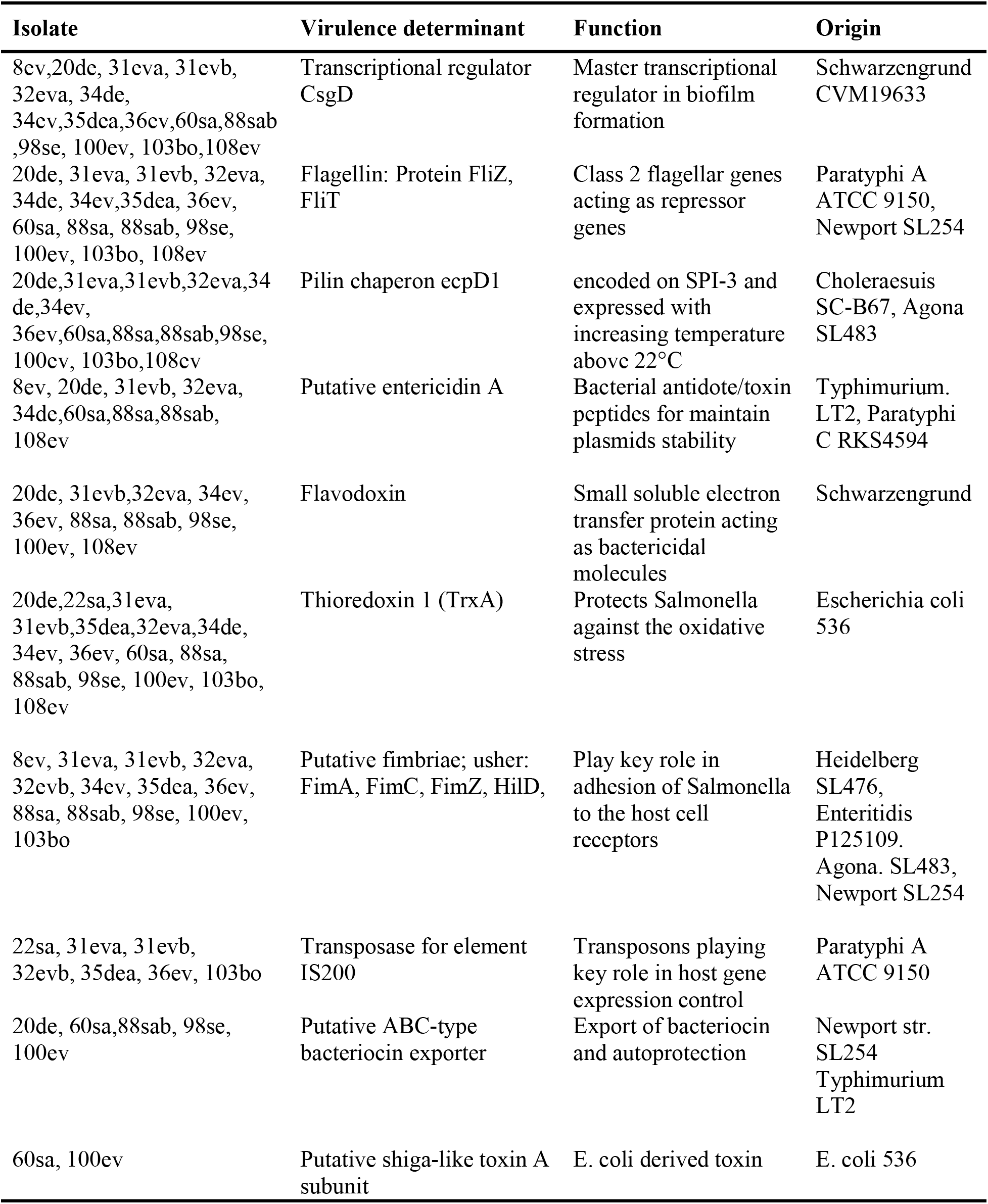
Distribution of adhesion factors, toxins and virulence plasmids.

**Table 8b:** Distribution of adhesion factors, and virulence plasmid.

| Isolates | Virulence determinant | Function | Origin |
| --- | --- | --- | --- |
| 8ev, 22sa, 34de | Col (PHAD28) plasmid | Harbor gene *qnrB* encoding resistance against fluoroquinolone | S. Hadar HAD28 |
| 8ev, 20de, 34de, 60sa | IncFII (S) and IncI1-1 ($\alpha$) plasmids | Harbor toxin SpvB and mediate resistance against aminoglycosides, tetracycline, chloramphenicol, sulfonamides, β-lactams, fluoroquinolones | Paratyphi C RKS4594 Typhimurium |
| 34de | IncH12 and IncH12A plasmids | Mediate resistance to aminoglycosides, tetracycline, chloramphenicol, sulfonamides, β-lactams, fluoroquinolones | Serratia mrescens K478 |
| 8ev, 20de, 34de 31eva, 32eva, 32evb, 35dea, 36ev, 88sab, 98se, 100ev | Proteins CuSA & ScsC | Regulate the intracellular excess of copper and promotes intramacrophage survival | Agona SL483, Typhimurium LT2 Newport SL254 |
| 8ev, 20de, 31eva, 31evb, 32evb, 34de, 34ev, 35dea, 36ev, 60sa, 88sa, 98se, 100ev, 103bo, 108ev 88sab | Putative Lys R family | Regulates many cellular processes (activators and repressors, N-terminal DNA binding domain) | Paratyphi A ATCC9150, Typhimurium LT2 |
| 34de, 60sa, 88sa, 88sab, 98se, 100ev | Virulence genes *terC*, *tehA*, *Terx*, *Terw* | Plasmid-borne genes coding for tellerium ion protein resistance | E. coli plasmid pJIE186-2 |
| 34de | *cvaC* and *mchF* genes | Microcin V genes for the synthesis of microcin | E. coli plasmid<sup>PCOLVCA 7V</sup> |

While 82% *Salmonella* isolates had the necessary virulence factors (SipABCD, SopBDE, SopE2, EnvEF, InvAE, Sii E, IagB, SptP, OrgB, PhoP, MisL, ShdA, rtn) to gain access into the host cells and to esblish infection, between 56 and 61% expressed adhesion factors (FimA, FimC, FimZ, HilD, ecpD1, CsgD, FliZ, FliT). The effectors responses (SopABDE, SpiC, SpvB, SseBCD, PipB2, SptP, SopE2, proteins Spa family) that mediate maturation, tracfficking process and replication within the SCV and control of the host immune were present in between 56 and 69% *Salmonella* isolates. Additionally, while 95% NTS carried H-NS, the master silencer of horizontal transfer genes, between 21 to 47% expressed heat shock and stress proteins, CuSA, ScsC, bacteriocin exporter.

Four types of plasmids (IncF plasmids, IncI plasmid, Col (PHAD28) plasmid and IncH plasmid) and four bacterial toxins (Flavodoxin, Putative Shiga-like toxin A subunit, Putative entericidin A and Thioredoxin 1) were detected in *Salmonella* isolates.

### Genotypic antibiotic resistance profile of *Salmonella* isolates

The total lengths of these genomes ranged from 4.5 to 4.8 Mb, with an average GC content of 51.7%, and the numbers of contigs ranged from 50 to 359 (Table S1). The numbers of protein-coding genes, ncRNAs, tRNAs, and CRISPR arrays rangedi from 3,246 to 5,881, 2 to 15, 13 to 71, and 2 to 3, respectively. The GenBank accession numbers are found in Table S1.

## Discussion

Despite a relatively moderate resistance shown to streptomycin (21.73%), the majority of *Salmonella* strains isolated at the Yaounde slaughterhouse were sensitive to tetracycline, chloramphenicol, and ampicillin (Table 1). These results corroborate the findings of previous studies that showed common resistance against streptomycin among *Salmonella* [6]. In contrast, 75% of European *Salmonella* isolates were resistant to tetracycline, chloramphenicol, ampicillin, and streptomycin [3]. Findings from the present study indicate that tetracycline, chloramphenicol and ampicillin are still effective antibiotics in Cameroon.

The presence of 13 % MDR *Salmonella* isolates constitute a serious health concern. Similarly, a study conducted in Ethiopia in 2012, reported MDR to ampicillin, streptomycin, tetracycline and chloramphenicol among *Salmonella* isolates [7]. In the present work, MDR salmonellae were isolated from beef at different processing steps. This observation supports the conclusions of previous studies that considered cattle as one of the main reservoirs of resistant *Salmonella* to humans (8). This calls for public awareness on the danger of eating raw or insufficiently cooked beef meat.

The detection of the respective resistance genes to streptomycin, chloramphenicol, tetracycline and ampicillin in the *Salmonella* isolates (Table 2) confirms their phenotypic antimicrobial resistance status described in Table 1. However, false negative cases were reported for each of the tested antibiotics. This mismatch between phenotypic and genotypic antimicrobial resistance was observed by previous authors who believed that some antimicrobial-resistant genes are “silent” in bacteria *in vitro* [9–12]. The “silent genes”, are antibiotic resistance genes that were detected in the susceptible *Salmonella* isolates but failed to be expressed phenotypically. The exhibition of the resistance phenotype depends on the mode and level of expression of the resistance gene that could be influenced by cultural or other environmental factors [9]. However, the reasons why these genes were not expressed phenotypically are yet to be elucidated. Based on information reported by previous studies [9–12], the reason could be due the absence or impairment of the promoter sequence required for their expression. Another reason could be the presence of negative regulators of resistance genes that migt have downregulated their expression in vitro. It could also be that the expression of these genes might have occurred but without the expression of the gene products in the isolates. It appears that many factors exist to control the expression of the resistance phenotypes. The detection of such genes in the *Salmonella* isolates may have important implications. These genes will not always be locked on the “silent mode”but there is a possibility that they can be activated thereby inducing phenotypic resistance in the host *Salmonella* [30]. Another implication of the “silent genes” is that the host organisms may become an important reservoir of antibiotic-resistance to susceptible or major food pathogens beyond the *Salmonella* species via horizontal gene transfer or other mechanisms [10].This explains why several scientists considered “silent genes” as a significant potential threat to the therapeutic efficacy of antibiotics [10–11]. Remarkably, all the three *Salmonella* strains that exhibited phenotypic multidrug resistance were all Poona serovars and harbored the respective resistance genes (Tables 2 & 3).

The 18 genes involved in multidrug resistance are basically divided into three groups based on their mode of actions [10]: (i) those involved in exporting drugs out of the bacterial cells (*sdiA, mdsA, mdsB, mdsC, E. coli soxS, E. coli soxR, acrB, acrD, golS, marA, patA* and *ramR*) constitute the majority; (ii) those that encode for reducing membrane permeability to drugs (*E.coli soxR, E. coli soxS* and *marA*) and (iii) those involved in antibiotic target alteration (*bacA, E. coli soxR, E. coli soxS* and *ramA*). However, *E.coli sox R, E. coli soxS* and *ramA* genes have an overlapping role and act at the same time in mediating drug efflux, reducing drug entry into cells or modifying the configuration of the antibiotic target (Data not shown).

All the genes involved in drug efflux are part of the resistance nodulation cell division (RND) efflux systems and only effectively perform their duty in synergy with TolC [11]. Moreover, there were also multi-efflux pumps such as *mdsA, mdsB, mdsC, acrA, acrB*, and *mdtk*, which generally work in synergy either with transcriptional activators (*E,coli soxS* and *E.coli soxR, ramA, marA*) or promotors such as *TolC* and *golS* [13]. The presence of *TolC* only in the phenotypically multidrug *Salmonella* isolates (**8ev**, **22sa** and **34de**) justifies the critical role it plays in synergy with RND efflux systems. TolC is member of the outer membrane efflux protein (OEP) family involved in exporting a range of antimicrobials and other harmful compounds such as metals via either funnel or switch mechanisms [12]. The *golS* gene promotes MdsABC complex to confer resistance against a variety of drugs and toxins and to confer virulence and pathogenicity potentials to *Salmonella* [14]. This justifies why the MdsABC complex was also found in all the isolates that had *golS* promoter.

The gene *sdiA*, a regulator for *AcraB*, a multi-drug resistance pump [15]detected in 100% of the isolates is as powerful as *E.coli soxS* and *E. coli soxR* in providing resistance against a vast arsenal of antibiotics including Rifamycin. The presence of *mdtK* and *qnrB*1 in our isolates is of paramount importance. These two genes synergistically offer *Salmonella* advantage to develop resistance via plasmid-mediated or efflux pump mechanism against fluoroquinolone considered highest priority drug [5,16]. However, resistance to fluoroquinolone could also be due to transcriptional activators such as mutated *ramR, soxS and marA* usually associated with the overexpression of *acrAB-TolC* efflux pump in *Salmonella* species [17].). Equally, the *E.coli soxS and E.coli soxR* genes probably borrowed from *E.coli* confer resistance against all the classes of antibiotics including fluoroquinolone.

The detection of *OXA* (OXA-1, *OXA-2* and *OXA-7*) genes in **8ev, 22sa and 34de** isolates, *CTX-M-14* gene in **8ev** and the gene *qnrB* in isolates **8ev**, **20de**, **22sa**, **34de**, and **60sa** is another area of great concern as 3^rd^ generation cephalosporin and fluoroquinolone are considered highest priority drugs [16]. Moreover, broad-spectrum cephalosporin-hydrolyzing and carbapenem-hydrolyzing activities of respectively, OXA-1 and OXA-2 were recently reported [18–19]. In general, these results call for an urgent investigation for antimicrobial resistance status of non-typhoidal *Salmonella* circulating in Cameroon.

Polymyxin is a bactericidal polypeptide, which disrupts lipid A subunit of the LPS outer membrane of Gram-negative bacteria causing cell lysis and their eventual irreversible death. *Salmonella* are among Gram-negative organisms that used to develop acquired resistance to polymyxins in opposition to *Proteus*, *Serratia*, etc, which are naturally resistant to this peptide molecule [20]. One of the key mechanisms adapted by the Gram-negative bacteria including *Salmonella* resides in the modification of the lipid A contained in the bacterial LPS outer membrane [19]. This may occur because of mutations in the PmrA/PmrB causing overexpression of LPS-modified genes. At the molecular level, lipid A misconfiguration involves the addition of 4-amino-L-arabinose (LAra4N) to phosphate groups within the lipid A thereby reducing the permeability of polymyxin into the bacterial cells via decrease in the net charge of the modified lipid A [21]. Moreover, the presence of mutations in pmrA only in isolates **8ev**, **22sa** and **34de** reinforces the multidrug resistance status of *Salmonella* Poona in this study. The resistance potential of *Salmonella* isolates to polymyxin in this study via the observed mutations in pmrA/pmrB unveils both clinical and veterinary importance due to their zoonotic nature [20].

All the *Salmonella* isolates (Table 5) have high probability (with a mean probability of 94%) to cause diseases in humans. Moreover, their belonging to a large pathogenic families (466-787) confirms the zoonotic status of nontyphoidal *Salmonella* and their ability to exhibit broad-host adaptation. It is not surprising to see serovar Enteritidis scoring the highest probability to cause disease in humans. This serovar has been ranked among the ten most important foodborne diseases by CDC and as the most prevalent foodborne pathogen in the world after Typhimirium based on its involvement in disease outbreaks [22]. It should be recalled that 50% of Enteritidis isolates in the current study belong to sequence type ST-11, which is incriminated to be the major cause of African iNTS as described by many authors [22]. However, though exceptionally multidrug resistant, serovar Poona appears to be the least human pathogen. This relative low pathogenicity to human may underline their animal host restriction. These findings suggest for adequate cooking of beef carcass and sensitization of some butchers who still have the habit to eat raw meat.

The *Salmonella* pathogenicity islands (SPIs) which represent powerful weapons for *Salmonella* species were described in Table 6. The constant presence of C63PI in all the isolates may explain its indispensability for survival as this is required for iron uptake (Table 7). Previous studies are in favor of the conservation of C63PI among *Salmonella* isolates [23–24]. Eleven SPIs including C63PI have been identified in our isolates. Five appear (SPI-1, SPI-2, SPI-3, SPI-4 and SPI-5) to play a more critical role in *Salmonella* infections though all the virulence factors are generally interconnected via a complex network of Type III secretion system. The remaining detected pathogenicity islands (SPI-8, SPI-9, SPI-13 and SPI-14) are involved in one way or the other in the regulation of the expression of other SPI genes or the associated effector proteins (Table 7). While SPI-1 is more involved in the initiating stage of *Salmonella* infection, SPI-2 is required for systemic infection and intracellular pathogenesis by facilitating replication of intracellular bacteria within Salmonella-containing vacuole (SCV) [21]. While SPI-3 is required for survival in macrophages and conferring ability to *Salmonella* to grow in the low-magnesium environment, SPI-4 is needed for intramacrophage survival, for toxin secretion and apoptosis [25]. Moreover, SPI-5 is involved in the induction of pro-inflammatory immune response with diarrhoea as one of the outcomes in cattle [26]. However, overlapping or complementary functions have been observed between SPI-1, SPI-2, SPI-3, SPI-4 and SPI-5. This observation was noted by previous authors [27]. In general, the outcomes of *Salmonella-host* interaction depend on the host, the serovar involved in the interaction and the strength of the immune system of the host. The observed variation in SPI profile among NTS in this study unveils their differential degree of virulence. Our findings corroborate the results of the study carried out on poultry isolates in Nigeria [24]. It should be noted that nine other pathogenicity islands have not been detected in our isolates likely due to loss as *Salmonella* organisms evolved in the course of time.

Each of the SPIs works in synergy with an arsenal of genes that code for different specialized effector proteins. These effectors are either secreted or translocated into the host cells to enable *Salmonella* manipulate the host key cellular functions such as signal transduction, membrane trafficking and immune responses. About 82% of *Salmonella* isolates in the present study were endowed with not all but the necessary virulence factors (SipA, SipC, SopB, SopD, SopE, SopE2, SipB, EnvE, EnvF, InvA, InvE, SipD, Sii E, IagB, SptP, OrgB, PhoP, MisL, ShdA, rtn) to gain access into the host cells in order to esblish infection. Interestingly, all the effectors described in this phase of the infection are all SPI-1 encoded, indicating its active role in the *Salmonella* entry process [28–29]. Putative fimbriae usher (FimA, FimC, FimZ, HilD), pilin chaperon (ecpD1, CsgD) and flagellin (proteins FliZ, FliT) collectively called adhesion factors are additional virulence determinants to ensure successful colonization and persistence of *Salmonella* in the hosts [23]. Moreover, the expression of SipA and SipC required for invasion efficiency and of SopE and SopE2 for internalization efficiency [28–29] reinforced the pathogenicity potentials of our isolates. Despite the absence of SseI, SseJ, SspH and SifA, the expression of other effectors including SopB, SopE, SopA, SopD, SpiC, SpvB, SseB, SseC, and SseD, PipB2, SptP, SopE2 and antigen presentation proteins SpaK, SpaN, SpaS, and SpaR could still enable *Salmonella* isolates to achieve the maturation and tracfficking process of the Salmonella-containing vacuole, to replicate within the SCV and to control the host immune responses. The absence of the aforementioned virulence factors in our isolates may underline the variation in the virulence potential of *Salmonella* species. Additionally, *Salmonella* isolates were also well armed with effectors such as heat shock protein, stress protein, proteins CuSA and ScsC, putative ABC-type bacteriocin exporter to successfully compete with other bacteria and to be able to control their growth environment [27, 30–31]. Apart from the invasion phase which, was SPI-1)dominated, other stages of *Salmonella* pathogenesis (survival and replication within SCV and macrophages and anti-inflammatory immune responses) involved the participation of other SPIs including SPI-2, SPI-3, etc. This explains probably why some virulence determinants such as SipA, SopB and SopD carried out more than one biological activities. Though, HilA, the leading regulator of SPI-1 [29], was not detected in the isolates, however, between 59 and 68% of NTS isolates possessed PhoP, HilD, and FliZ, three important transcriptional regulators while 95% carried H-NS, the master silencer of horizontal transfer genes [27, 32] to regulate the secretion or translocation of effector proteins during each step of *Salmonella* infection [33]. The absence of HilA was very likely due to sequencing error in the coding regions of their respective coding genes given the presence of other key transcriptional regulators including PhoP, HilD, and FliZ.

Unlike other virulence factors, virulence plasmids were not widely distributed among NTS in this study. IncF plasmids known as crucial virulence plasmids in *Salmonella* [12] were carried by *Salmonella* isolates that harbored the gene *qnrB*. Similarly, recent study in China indicated that *Salmonella* isolates harboring IncF plasmids were all fluoroquinolone-resistant [34]. Though IncI1 plasmids seem to be widely distributed in *Salmonella* [12, 35], only close to 23% NTS at the Yaounde abattoir were positive for their presence. Even though there was no evidence of serovar preference, Col (PHAD28) plasmid was only present in serovar Poona (**8ev**, **22sa** and **34de**), which were all multidrug resistant. Their multiple resistance potential corroborates with previous study indicating that most *Salmonella* harboring this plasmid were multiple antibiotic resistant [12]. On the other hand, IncH plasmids were only expressed in isolate **34de**. Previous study reported the presence of such plasmids in *Salmonella* that carried on their genomes the genes *cat, qnr, strAB*, TEM-1, and *tet* [12]. This corroborates with our findings indicating the presence of the same genes in isolate **34de**. Remarkably, all the plasmids carried by *Salmonella* in this study are mobilizable plasmids. Mobilizable plasmids in *Salmonella* are known to promote multiple drug resistance [12]. One of the major limitations to this study was the loss of some information during sequencing. However, despite this limitation, the present study unveils the multidrug resistance, pathogenicity and virulence potentials of *Salmonella* isolates at the Yaounde abattoir.

### Conclusion

Results revealed the public health importance of *Salmonella* isolates at the Yaounde abattoir. Phenotypically, while around 26% of *Salmonella* isolates exhibited resistance to streptomycin, 13 % were MDR. Genotypically, all the *Salmonella* isolates possessed high MDR potentials and harbored *OXA-1*, *OXA-2*, *OXA-7*, *CTX-M-14* and *qnrB1* genes conferring resistance to 3^rd^ generation cephalosporin and fluoroquinolone, respectively. More than 31% of the isolates exhibited resistance potential to polymyxin, considered as critically important drug. Additionally, close to 80% of NTS isolates harbored ‘’silent resistant genes’’ which may act as reservoir of resistance to foodborne bacteria. Detected plasmids in this study seem to promote MDR in *Salmonella* and should be integrated in the antibiotic surveillance programme. *Salmonella* isolates also exhibited a high degree of pathogenicity and virulence to establish infection in their hosts including humans. Key virulence factors, SPI-1, SPI-2, SPI-3, SPI-4 and SPI-5 including important effector proteins were present in many isolates. The Whole genome sequencing technique is an important tool in detecting MDR and pathogenicity potentials of NTS. This calls for the prudent use of antibiotics, and constant monitoring of AMR of NTS in Cameroon. In perspective, phenotypic resistance profile against the antibiotics for which the resistance genes were detected is required to establish the overall multidrug resistance status of our *Salmonella* isolates.

## Materials and Methods

### Bacterial isolates

This work was approved and an ethical clearance was granted by the Cameroon Bioethics Initiative Review and Consultation Committee (CAMBIN, Ref CBI/ 406/ ERCC/CAMBIN). NTS were isolated from live cattle (n=1), beef carcasses (n=19), processing environments (n=2), and butchers’ hands (n=1) at the Yaounde abattoir following ISO 6579 [36] between December 2014 and November 2015. All *Salmonella* isolates were confirmed using API-20 E kit (BioMérieux, France) and a qualitative real-time PCR assay [37].

### Antibiotic susceptibility tests

*Salmonella* concentration (1.5×10^8^ CFU/ml) and those of controls (*Escherichia coli* ATCC 25922 and *Staphylococcus aureus* ATCC43300) were spread onto the surface of Mueller-Hinton agar to which the antibiotic disks were placed and incubated for 18 to 24 hours. The diameter of the zones of inhibition around each antibiotic disk were measured with a ruler and recorded to the nearest millimeter and isolates were classified as resistant, susceptible, intermediate [12]. The antibiotics used were ampicillin (AMP) 10 μg, chloramphenicol (C) 30 μg, tetracycline (TE) 30 μg, and streptomycin (STR) 25 μg.

### DNA extraction

Total DNA was extracted from *Salmonella* overnight culture using Quick-DNA™ Miniprep Plus Kit (Zymo Research, Irvine, CA) following the manufacturer’s instructions. The purified DNA was quantified using a NanoDrop 2000c spectrophotometer (ThermoFisher Scientific, USA) and stored at −20°C until use.

### Library preparation and sequencing

Paired-end libraries were constructed with 0.2 ng/μl of purified DNA using the Nextera XT DNA Library Prep Kit as recommended by the manufacturer (Illumina, San Diego, CA, USA) and were quantified using Qubit fluorometer (ThermoFisher Scientific, USA). WGS was performed on Illumina NextSeq platform using NextSeq 500/550 high-output kit v2 (300 cycles) at Murdoch University (Australia) and on Illumina Miseq platform using pair-ended MiSeq reagent v3 kit (2 × 201 bp) at the BecA-ILRI Hub (Nairobi, Kenya) following the manufacturers’ guidelines.

### Resistance genes, pathogenicity, and virulence factors

The quality of the raw sequences were checked with FASTQC and trimmed using Trimmomatic 2.6 at Q score below 20. The trimmed data were assembled using SPAdes version 3.11, and genomes were annotated with the NCBI Prokaryotic Genome Annotation Pipeline [39]. The antibiotic resistance genes (ARG) in the assembled genomes were identified by BLAST search against the reference ARG sequences from ResFinder 3.0 [40] and CARD 2017 [41]. The pathogenicity of NTS was predicted using PathogenFinder 1.1 [42]. The *Salmonella* pathogenicity islands were detected using SPIFinder 1.0 [43]. Acquired virulence genes were detected by uploading the raw reads into VirulenceFinder 2.0 [44] while virulence plasmid was assessed using PlasmidFinder 2.1 [45].

## Supporting information

The Salmonella genome sequence data reported here have been deposited into NCBI GenBank under BioProject PRJNA432303. The GenBank accession no of the *Salmonella* genomes are in Table S1

## Acknowledgements

We thank the University of Udine, Italy for providing the Qiagen extraction kit and Real-Time PCR reagents. We also thank Profs Mary Barton and Darren Trott (University of Adelaide, Australia), Prof Sam Abraham and Dr Mark O’Dea (Murdoch University, Australia), for sequencing facilities. Finally, we are grateful to Joyce, and Jean Baka for the Bioinformatics support.

## Author Contributions

**Conceptualization**: Chelea Matchawe, Edi Piasentier, Roger Pelle.

**Data curation**: Chelea Matchawe

**Formal analysis**: Chelea Matchawe.

**Funding acquisition**: Chelea Matchawe

**Investigation**: Chelea Matchawe.

**Methodology**: Chelea Matchawe, Eunice Matchawe, Martina Kyallo, Isaac Njaci, Dedan Githae, John Juma, Patrice Bonny, Nkeunen Gerard.

**Project administration**: Roger Pelle.

**Resources**: Edi Piasentier, Martina Kyallo, Eunice Machuka.

**Supervision**: Lucy M. Ndip, Edi Piasentier, Roger Pelle.

**Validation**: Chelea Matchawe.

**Visualization**: Chelea Matchawe.

**Writing** – original draft: Chelea Matchawe.

**Writing** – review & editing: Patrice Bonny, Gerard Nkeunen, Seraphine Esemu, Mohamadou Bawe, Bonglaisin J. Nsawir, Martina Kyallo, Eunice Machuka, Roger Pelle

